# Venomous Maggots? A first exploration of the toxin arsenal of larval stages of the horse fly *Tabanus autumnalis*

**DOI:** 10.1101/2025.05.06.652366

**Authors:** Jonas Krämer, Ludwig Dersch, Andreas Vilcinskas, Tim Lüddecke

## Abstract

As a rapidly evolving trait, animal venom exhibits compositional variation even at the intraspecific level between populations, ontogenetic stages and sexes. In extreme cases venom is used for different functional purposes across ontogenetic stages. This phenomenon occurs for example in case of horse flies (Tabanidae), which utilize venom for predation during the larval stage and for blood feeding in the adult stage. While the venom composition and activity of adult tabanids has been investigated for some species, nothing is known about the venom compounds of tabanid larvae. With the current study we provide first insights into the venom composition of larval stages of a tabanid. Besides a plethora of hydrolyzing enzymes, we find the venom of *T. autumnalis* larvae dominated by peptide toxins. Some of these peptide toxins could be annotated as putative neurotoxins and cytolytic peptides and these compounds likely are the main drivers of fast prey incapacitation. Previous studies identified the main functional constituents utilized by tabanid imagines for blood feeding to be protease inhibitors affecting vasoconstriction and thrombin to promote blood feeding. Interestingly we find highly similar putative protease inhibitors in the larval venom, though with relatively low expression levels. This provides a first indication for a shifted expression profile between the venom of tabanid life stages to fulfill both the predatory needs of the larvae and the blood lust of female imagines.

## 1. Introduction

Animal venoms are highly complex mixtures mainly composed of proteinaceous components, which are injected into target organisms for the purposes of predation, host-parasite interaction, defense and intraspecific competition (Schendel et al., 2019). The composition of animal venom is highly variable and subject to various evolutionary pressures and environmental cues (Casewell et al., 2013). Consequently, venom variation is frequently observed at the intraspecific level and can occur between life-history stages, sexes or different populations (Chang et al., 2015; Daltry et al., 1996; Surm and Moran, 2021a). In the case of predatory venoms, such intraspecific differences might be driven mostly by differences in prey preferences. However, in some cases an alteration of the venom composition coincides with a purpose shift. For instance, some arachnids exhibit sex-specific differences in their venom composition that most likely are not related with different prey preferences (e. g. Binford et al., 2016; Krämer et al., 2022). Instead, besides predation, in the adult stage venom is additionally used in a sex-related context (Krämer et al., 2022; Zobel-Thropp et al., 2018) or for the purpose of defense (de Oliveira et al., 2005; Herzig et al., 2020). An extreme case of intraspecific venom-variation can be expected for holometabolous insects as metamorphosis promotes the specialization towards a secondary trophic niche (Ten Brink et al., 2019). However, the extent of venom modulation within the life cycle of an insect has not been explored yet.

Within insects, venom usage has evolved at least 14 times independently resulting in various venom delivery systems (Andrew A. Walker et al., 2018) and recent studies provided valuable insights into the venom compositions of several hitherto unexplored insect groups (e. g. Drukewitz et al., 2018; Fischer et al., 2024, 2023; Krämer et al., 2024; Qu et al., 2023). Insect venom usage can be restricted either to the adult stage (e. g. hymenopterans), the larval stage (e. g. neuropterans) or it occurs across developmental stages. The latter is described above all for dipterans. For many dipteran families known to utilize venom in the adult-stage like Asilidae, Rhagionidae, or Tabanidae venom usage is also postulated for the larvae that are described as voracious predators (Schmidt, 1982; von Reumont et al., 2014). However, not a single larval venom compound has been identified yet. Of the venomous dipterans, tabanids are a promising model to study the extent of intraspecific venom variation. This is because tabanids utilize venom for completely different purposes during their life cycle. While the imagines use venom to facilitate blood feeding, larval tabanids seem to possess a highly effective venom to subdue their prey. The latter is emphasized by two case reports, the first one describing the predation of froglets by tabanid larvae (Jackman et al., 1983) and the second one reporting painful bites caused by tabanid larvae to humans working as rice harvesters (Otsuru and Ogawa, 1959).

With respect to the toxin arsenal employed by adult tabanids for blood feeding, some studies investigated the venom composition of different tabanid species utilizing transcriptomics and proteomics (Ribeiro et al., 2015; Xu et al., 2008). In addition, the activity of salivary gland extracts and purified venom compounds were tested mainly in the context of blood feeding (Kazimírová et al., 2001; Ma et al., 2009; Rajská et al., 2007; Xu et al., 2008). In this regard, the best characterized toxins of adult tabanids that interfere with hemostatic processes are on the one hand vasodilatory peptides (e. g. Takáč et al., 2006a) and on the other hand peptides inhibiting thrombin a major factor of the coagulation cascade (Xu et al., 2008). In addition, some immune-repressive peptides have been functionally characterized that exhibit for instance anti-inflammatory activities (e. g. Yan et al., 2008). In contrast to the imagines, the venom composition of larval tabanids remains completely unexplored.

Here, we aim to address this research gap by providing the first proteo-transcriptomic analysis of the venom composition of a tabanid larvae. This will provide first insights regarding the major constituents of larval tabanid venom and allow for a first assessment regarding the differences to the venom of tabanid imagines.

## 2. Material and methods

### 2.1. Collection and rearing of animals

Larval stages of *Tabanus autumnalis* were collected in the Rieselfelder, Münster, North-Rhine Westphalia, Germany. The larvae were collected from below the turf by utilizing a hand rake similar as described in (Philip, 1928). For rearing, each collected specimen was placed into a 15ml Falcon^™^ tube, equipped with a moistened cotton pad. To ensure sufficient air supply the lids of the tubes were punctured. Once per week the larvae were fed with a pinky maggot (*Lucilia sericata*).

### 2.2. Proteomics workflow

To trigger the release of venom, tabanid larvae were stimulated with electricity in the anterior section of their abdomen by using a self-made electrical forceps coupled to a Promed tens device (Promed GmbH, Farchant, Germany; pulse width 200–250 ms, pulse rate 60–130 Hz, voltage between 8 and 12 V). The exuding venom was collected with pulled glass capillaries produced with the pipette puller P-2000 (Sutter instrument Novato, CA 94949, USA) and transferred into 1.5 ml tubes with 20µl Millipore water. Venom samples were stored at - 20°C until further use.

For each of the collected venom samples MALDI-TOF mass fingerprints were generated with a Bruker ultrafleXtreme TOF/TOF mass spectrometer (Bruker Daltonik GmbH, Bremen, Germany). For this purpose, the 1µl of venom stock solution was diluted 1/10 and 1/20. The diluted samples were spotted in triplicates of 0.3µl onto a MALDI-TOF sample plate and mixed on plate with an equal volume of 10 mg/ml 2.5-dihydroxybenzoic acid (Sigma Aldrich, Steinheim, Germany) matrix, dis-solved in 50% acetonitrile/0.05 % TFA. For an optimal crystallization of matrix salts, samples were blow-dried with a hairdryer. The spectra were generated in reflectron positive mode in mass ranges of m/z 800 to 4500 and 3000 to 10 000. For external calibration, a mixture containing proctolin ([M+H]+, 649.3), Drm-sNPF-212-19, ([M+H]+, 974.5), Pea-FMRFa-12 ([M+H]+, 1009.5), Lom-PVK ([M+H]+, 1104.6), Masallatotropin ([M+H]+, 1486.7), Drm-IPNa ([M+H]+, 1653.9), Pea-SKN ([M+H]+, 2010.9), and glucagon ([M+H]+, 3481.6) was used. Ion signals were identified by using the peak detection algorithm SNAP from the flexAnalysis 3.4 software package. In addition, each spectrum was manually checked to ensure that the monoisotopic peaks were correctly identified.

To obtain fragment spectra of venom compounds from larval tabanids both a top-down and a bottom-up approach were used to analyze venom samples from two specimens of *Tabanus autumnalis*. Protein concentrations of venom samples were determined with a Direct DetectTM Spectrometer (Merck, Germany) and bottom-up proteomics was performed with 20 µg of crude venom per specimen and top-down proteomics with 10µg of crude venom per specimen. All samples were denatured in Urea buffer (8M urea/ 50mM triethylammonium bicarbonate buffer, final concentration of urea (7M)) followed by reduction and alkylation with dithiothreitol and chloracetamide. For those samples used for bottom-up, additionally a digestion was performed with trypsin (Sigma-Aldrich, USA) applying an enzyme/substrate ratio of 1/75. Desalting and removal of urea was achieved with poly (styrene divinylbenzene) reverse phase (RP)-StageTip purification. Mass spectra were generated with a Q-Exactive Plus (Thermo Fisher Scientific) mass spectrometer coupled to an EASY nanoLC 1000 UPLC system (Thermo Fisher Scientific, Bremen, Germany). For UPLC in-house packed RPC18 columns with a length of 50 cm were used (fused silica tube with ID 50 µm +/-3 µm, OD 150 µm; Reprosil 1.9 µm, pore diameter 60 A°; Dr. Maisch GmbH, Ammerbuch-Entringen, Germany). The UPLC upstream separation of venom compounds/tryptic peptides was performed with a binary buffer system (A: 0.1% formic acid (FA), B: 80% acetonitrile, 0.1% FA): linear gradient from 2 to 62% in 110 min, 62–75% in 30 min, and final washing from 75 to 95% in 6 min (flow rate 250 nL/min). Re-equilibration was performed with 4% B for 4 min. To obtain fragment spectra with the Q-Exactive Plus, HCD fragmentations were performed for the 10 most abundant ion signals from each survey scan in a mass range of m/z 300–3000. The resolution for full MS1 acquisition was set to 70,000 with automatic gain control target (AGC target) at 3 * 10^6^ and a maximum injection time of 80 ms. The run was performed at a resolution of 35,000, AGC target at 3 * 10^6^, a maximum injection time of 240 ms, and 28 eV normalized collision energy; dynamic exclusion was set to 25 s.

### 2.3. Transcriptomics workflow

The same two larval specimens from which venom was extracted for top-down and bottom-up proteomics were used to extract RNA for transcriptomics. To obtain the venom glands (salivary glands) the animals were anesthetized by placing them on ice for 10 minutes and dissected in ice-cold phosphate-buffered saline (PBS). The venom glands and for one of the specimens additionally parts of the midgut were carefully separated from the surrounding fat body and each tissue sample was transferred into 1ml TRIzol (Thermo Fisher Scientific, Darmstadt, Germany) followed by homogenization with scissors and sonic finger (15 s). Then RNA-extraction was performed according to the standard TRIzol user guide and RNA-concentration was with Qubit (Thermo Fisher Scientific, Waltham, Massachusetts USA). Library preparation was achieved for 1µg of total RNA using the Illumina® TruSeq® stranded RNA sample preparation Kit. The libraries were validated and quantified with an Agilent 2100 Bioanalyzer and paired-end sequencing was performed with an Illumina TruSeq PE Cluster Kit v3 and an Illumina TruSeq SBS Kit v3—HS on an Illumina HiSeq 4000 sequencer. For quality trimming and adapter removal of raw data, Trimmomatic 0.3.8 (Bolger et al., 2014) was used and transcriptome de novo assembly was performed with Trinity v2.8.5 (Grabherr et al., 2011), to generate one single assembly based on all raw data applying default settings. Afterwards getorf from EMBOSS v. 6.6.0.0 (Rice et al., 2000) was used to perform a six-frame translation and to extract open reading frames. Finally, the transcript expression levels were estimated with kallisto v. 0.46.1 (Bray et al., 2016) for each raw dataset.

### 2.4. Identification of venom precursors

The first step to identify the venom compounds of larval tabanids was matching the proteomic data against a database based on the generated transcriptome data using the software PEAKS 12.5 (PEAKS Studio 12.5; BSI, Toronto, ON, Canada). For both, the top-down and the bottom-up data, combined PEAKS searches (comprising the data of both larval specimens) were performed against the six-frame translation of the transcriptomic data. The searches were performed with a parent error mass tolerance of 10 ppm and a fragment mass error tolerance of 0.05 Da. Carbamidomethylation was set as fixed post-translational modification (PTM), whereas acetylation (N-term), amidation, carboxymethyl, half of a disulfide bridge, oxidation (at M) and pyro-glu from E/Q were considered as variable PTMs. Enzyme mode was set to ‘None’ with ‘Unspecific’ digest mode for the top-down run and for the bottom-up analysis these parameters were set to ‘Trypsin’ and ‘Semi-Specific’ respectively. For all venom precursors with proteomic coverage SignalP v. 6 (Teufel et al., 2022) was used to check for the presence of a signal peptide with slow–sequential mode and eukarya as parameters. Next, a filtering step was performed by removing all precursors without signal peptide and by setting the PEAKS parameters ‘Coverage’ and ‘10*lg(P)’ to minimum values of 7% and 30 respectively. Finally, the PEAKS-results were manually inspected to identify untypical propeptide cleavage sites and to assess the general coverage of proteomic and transcriptomic data.

### 2.5. Annotation of venom precursors and grouping into protein families

Venom precursors were analyzed with InterPro v. 5.72-103.0 (Blum et al., 2021) and searched against different subsets of the UniProt database (The UniProt Consortium, 2021) using BLAST 2.12.0 (Camacho et al., 2009). All venom precursors for which an InterPro-ID was allocated were annotated based on the corresponding InterPro-family entry or InterPro-description. Venom precursors without informative InterPro-results were classified based on BLAST-searches against precursors from the UniProt-database. For this purpose, separate BLAST searches were performed against the reviewed subset of the Toxprot database (access date: 14.01.2025, search term:’(taxonomy_id:33208) AND ((cc_tissue_specificity:venom) OR (keyword:KW-0800)) AND (reviewed:true)’) and against all reviewed Metazoan precursors deposited in the UniProt database (access date: 14.01.2025, search term: ‘(taxonomy_id:33208) AND (reviewed:true)’). Additional BLAST-searches were performed against these database subsets but also including the unreviewed entries. For each BLAST-search the ‘best’ hit was selected among the 10 hits with the lowest E-value considering bitscore, percentage identity, query coverage and BLASThit coverage. In addition, a bitscore threshold was used to filter low quality BLAST-hits. Finally, of the BLASThits with the different subsets of the UniProt-database the most informative was selected for functional annotation. For all peptide toxin precursors, additionally the alignments with their corresponding ‘best’ BLASThit were inspected to verify if the alignment covers the putative mature peptide.

For grouping the identified venom precursors into isoform families BLAST was used to assess the similarity between all identified venom precursors. For precursor sequences with a similarity above 70%, alignments were manually inspected and based on this, precursors were grouped into isoform families.

## 3. Results

The database used for matching venom proteomes of tabanid larvae, was constructed based on transcriptomes from salivary glands of two larval specimens and midgut tissue from one of these specimens. Sequencing RNA extracted from the salivary glands produced 19,497,599 (16,704,588 after trimming and quality control) and 22,975,878 (20,613,282 trimming and quality control) raw read pairs respectively, whereas 16,498,808 (13,789,298 after trimming and quality control) raw read pairs were obtained for RNA from the midgut tissue. These raw data were assembled into a single transcriptome assembly comprising 93,510 contigs and assessing the completeness with BUSCO yielded 90.3% complete BUSCOs (C: 90.3% [S: 50.6%, D: 39.7%], F: 1.7%, M: 8.0%, n: 1367).

Matching the venom proteomes against the transcriptome database enabled the identification of 475 venom precursors after quality filtering. According to our thresholds applied for similarity-based functional annotation, nearly half of the identified venom precursors could not be annotated and were classified based on

Cysteine-content and sequence length of the assumed mature peptide as uncharacterized linear peptides (∼22%), uncharacterized Cysteine-containing peptides (∼15%), uncharacterized Cysteine-containing proteins (∼11%) and uncharacterized proteins (less than 1%) (Figure 1). Of those precursors that could be functionally annotated, most were classified as hydrolyzing enzymes mainly comprising putative peptidases (∼15%) and to lesser proportions putative triacylglycerol lipases (∼3%), putative phospholipases A2 (∼1%), putative hyaluronidases (less than 1%) and putative chitinases (∼1%) (Figure 1A). The putative peptidases are not only abundant in terms of precursor count but also account for approximately one quarter of the total expression level of all venom precursors (Figure 1B). Besides hydrolyzing enzymes, some other classes of proteins were discovered as part of the venom composition comprising putative peptidase inhibitors (∼7%), odorant-binding proteins (∼2%) and proteins with CRISP-domain (∼1%).

**Figure 1.**
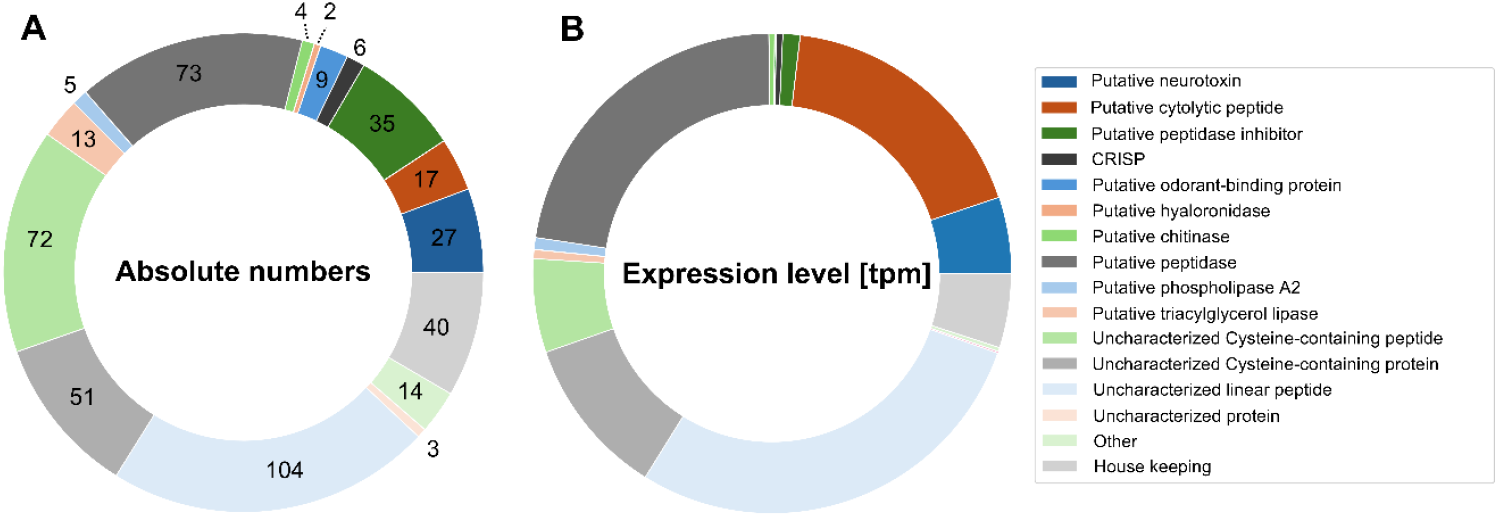
Venom composition of *T. autumnalis* revealed by a combined proteo-transcriptomic analysis. Venom compounds were classified according to their functional annotation based on similarity to known compounds from the uniport database **A**) absolute numbers of venom precursors identified per functional class **B**) Summed up expression levels (transcripts per million) per functional class of identified venom precursors.

A distinctive feature of the venom composition of larval stages of *T. autumnalis* is the high amount of peptide precursors which dominate the venom composition in terms of precursor count and contribution to the total expression level. Regarding their functional role in the venom, a fraction of these peptides could be annotated as putative neurotoxins (∼6%) and another fraction as putative cytolytic peptides (∼4%). Some of the identified neurotoxin precursors (U-Tabanustoxins-Ta1-3) exhibit mature peptides without Cysteines that show similarity to neurotoxins from ant venom from the Formicidae venom clade family 1, 3 and 5. Interestingly, the tabanid peptide toxins also exhibit the long pro-peptide region which is a characteristic feature also present in the corresponding BLAST-hits. U-Tabanustoxin-Ta2a is an outlier as this peptide precursor shows a higher similarity to antimicrobial peptides from the Mastoparan subfamily. Other putative tabanid neurotoxins comprise mature peptides with a six-Cysteine scaffold, showing similarity to conotoxins from the Conotoxin O1 superfamily on the one hand (U-Tabanustoxins-Ta7-9) and to asilidins from the Asilidin-1+12 families on the other hand (U-Tabanustoxins-Ta10-13). The remaining putative peptide neurotoxins contain 2 - 4 Cysteines and exhibit BLAST-hits with neurotoxins identified in scorpion and ant venom (U-Tabanustoxin-Ta14-16). Finally, two putative neurotoxin precursors (U-Tabanustoxin-Ta17+18) are processed into larger proteins containing 8 Cysteines and these show BLAST-hits to weakly annotated toxins from the Scoloptoxin-16 family.

Those peptides annotated as AMPs comprise some of the venom precursors with the highest overall expression levels. The largest fraction of these AMP-like precursors shows similarity to the cecropin-family, an AMP-family originally identified in moth hemolymph. It is noteworthy that most of these cecropin-like precursors (Cecropin-like-peptide-Ta1a-c) are not up-regulated in the venom-gland but show similar expression levels as in the non-venom producing midgut tissue. The one exception is U-Tabanustoxin-Ta5 which is a precursor highly expressed in the venom glands. The mature peptide processed from this precursor is most derived from all other identified cecropin-like peptides but exhibits a similar N-terminus compared to cecropin A from *Bombyx mori* hemolymph. Besides the cecropin-like precursors, one of the two isoforms of U-Tabanustoxin-Ta4 shows similarity to Moricin-1 another AMP identified for *Bombyx mori* with confirmed antimicrobial activities. For some of the tabanid venom precursors (U-Tabanustoxin-Ta2a, U-Tabanustoxin-Ta6), the best BLAST-hits are with AMPs identified in the venom of hymenopterans.

## 4. Discussion

Across the diversity of venomous animals, tabanids represent an extraordinary case with predatory larvae utilizing venom for prey incapacitation and parasitizing imagines that use it to facilitate blood feeding. So far, only the venom of adult tabanids has been analyzed in terms of composition and bioactivity and for the larval stages not a single venom compound has been identified. With the current study we explore the venom composition of a larval tabanid for the first time. Based on the identified venom compounds, we will discuss how the larval venom might fulfill its prey incapacitating function. Finally, by comparing our findings to existing knowledge on the venom of tabanid imagines, we will derive hypotheses on how tabanids can utilize venom for two highly deviating functions.

### 4.1. Major biological functions of T. autumnalis venom

The most noticeable characteristic of the venom composition of *T. autumnalis* is the high number of precursors that are processed into peptide toxins, Especially the high abundance of linear venom peptides without Cysteines is noticeable with similar abundances of such peptides mostly described for venom compositions of spiders belonging to the RTA-clade (Kuhn-Nentwig et al., 2021). Another distinctive feature of the venom composition of *T. autumnalis* is the high amount of venom precursors, whose sequence deviation from known compounds is too pronounced for functional annotation. Such high abundances of unclassifiable venom compounds are frequently discovered in venoms of taxonomic groups neglected by venom research. Also, for other dipterans like robber flies, a large fraction of venom compounds could not be annotated due to high degrees of sequence disparity from known proteins (Drukewitz et al., 2018).

#### 4.1.1. Neurotoxins facilitate prey capture

Some of the identified peptide toxins show similarity to neurotoxins identified in the venom of other invertebrates. Generally, neurotoxins found in animal venom are mostly Cysteine-containing peptides that bind to various types of ion channels, thereby interfering with signal transmission which among various other effects can cause rapid paralysis (Fry et al., 2009). Hence neurotoxins are considered a key component for fast prey-incapacitation, which is one of the main purposes of predatory venom.

Most of the putative neurotoxins identified in the venom of *T. autumnalis* contain six Cysteines and probably exhibit an ICK-fold, which is one of the most common folds recruited into animal venom (Undheim et al., 2016). Alongside BLAST-hits with Conotoxins of unknown function (U-Tabanustoxin-Ta7-Ta9), the most meaningful BLAST-hits were found for U-Tabanustoxin-Ta10-Ta13, which show similarity to putative neurotoxins of the Asilidin-1 (U-Asilidin(1)-Mar1a and U-Asilidin(1)-Dg12) and Asilidin-12 family (Asilidin(12)-Dg3b), all discovered in the venom of robber flies (Asilidae) (Drukewitz et al., 2018; Andrew A Walker et al., 2018). Robber flies are another group of dipterans that utilize venom for prey capture, but in this case both the larvae and the imagines are described as predators. Especially the adult robber flies are known to utilize potent venom to overpower even well-defended prey. As with tabanids, only the venom of the imagines has been studied in asilids, comprising proteo-transcriptomic analyses performed for a few species (Drukewitz et al., 2018; Andrew A Walker et al., 2018). Therefore, all BLAST-hits found with asilidins are venom compounds identified for adult robber flies. For all of these BLAST-hits paralyzing effects on insects have been described (Drukewitz et al., 2018; Jin et al., 2020), though the specific effects on ion-channels could only be revealed for U-Asilidin(12)-Dg3b. This toxin affects the ion channels Kv11.1/KCNH2/ERG1 by modulation of the voltage-dependence of the channel activation (Jin et al., 2020).

Interestingly, a large fraction of the putative neurotoxins identified in the venom of *T. autumnalis* are peptides without Cysteines (U-Tabanustoxin-Ta1-3). These were classified as neurotoxins based on similarity to putative neurotoxins identified in the venom of ants (Robinson et al., 2024; Touchard et al., 2018). These ant-neurotoxins also lack Cysteines and cause paralyzing effects to blow flies upon injection (Barassé et al., 2023; Robinson et al., 2024), although specific activities on ion channels have not been confirmed yet. A noteworthy feature of the precursor sequences of these ant-neurotoxins is the presence of a long propeptide, rich in glutamic acid and alanine residues, which is cleaved at an undescribed cleavage site (EA). U-Tabanustoxin-Ta1-Ta3 show a very similar pro-peptide, and the resulting mature peptides are supported by coverage of proteomic data.

#### 4.1.2. Potentiating effects via cytolytic toxins

Besides putative neurotoxins, some of the venom peptides discovered for *T. autumnalis* show similarity to linear cytolytic peptides without Cysteines. Such peptides often act by disrupting membranes, and their specificity depends *inter alia* on their net charge, amphipathicity, hydrophobicity and helix propensity (Rádis-Baptista, 2021). Besides affecting microbes, these compounds can also act on mammalian cell membranes and exert cytotoxic effects. In this case a direct functional role in envenomation of prey is assumed as the membrane-disrupting peptides act as spreading factors thereby potentiating the effectiveness of other toxins (Kuhn-Nentwig, 2003; Kuhn-Nentwig et al., 2019).

Some of the AMP-like precursors (U-Tabanustoxin-Ta4a-Ta6) are clearly up-regulated in the venom glands of *T. autumnalis*. Of these, U-Tabanustoxin-Ta4a and U-Tabanustoxin-Ta5 were annotated based on similarity to AMPs from *Bombyx mori* (Moricin-1 and Cecropin-A) (Hara and Yamakawa, 1995; Yamano et al., 1994). It is noteworthy that besides U-Tabanustoxin-Ta5 several other precursors (Cecropin-like-peptide-Ta1a-Cecropin-like-peptide-Ta1c) with similarity to insect cecropins were identified. However, U-Tabanustoxin-Ta5 is the only cecropin-like peptide highly expressed in the venom glands and relative to the other cecropin-like peptides the sequence of U-Tabanustoxin-Ta5 is most deviating from cecropins. This might be an indication of a specificity-shift from antimicrobial towards metazoan cell membranes. Cytotoxic effects can also be assumed for U-Tabanustoxin-Ta6, based on its similarity to M-poneritoxin a linear peptide from ant venom for which insecticidal and hemolytic activities have been demonstrated, besides broad activities against microbes (Orivel et al., 2001).

#### 4.1.3. Predigestive role of enzymatic components

Maggots in general cannot ingest large food particles and hence rely on extra-oral digestion (EOD) for feeding, which was first demonstrated in 1931 (Hobson, 1931). For some maggots it was shown that the needed hydrolyzing enzymes for EOD are at least partially stemming from the salivary glands (Valachova et al., 2014). For this reason, it is not surprising that the venom glands (salivary glands) of *T. autumnalis* are enriched with hydrolyzing enzymes comprising hyaluronidases, chitinases, triaglycerol lipases, phospholipases and various peptidases. The putative main function of these enzymes is to liquify prey tissue thereby enabling EOD. Interestingly, in arthropods the primary production site of digestive enzymes is the midgut (Cohen, 1995). Some venomous animals like arachnids still inject hydrolyzing enzymes from the midgut after utilizing venom for prey incapacitation (Fuzita et al., 2016). Tabanid larvae are not the first example of venom used for predigestion, as hydrolyzing enzymes have also been found in the venom glands of other lineages, such as assassin bugs (Cantón and Bonning, 2020).

### 4.2. Switches in venom chemistry coincide with life history stage-bound feeding ecology

The different functional context in which tabanid larvae and imagines utilize their venom, raises the question to which extent the venom composition must be altered to fulfill both functional roles. In this regard two extreme hypotheses are plausible.

The first hypothesis is that the venom composition is similar across the life stages. This implies that either some of the toxins have a dual function or that unneeded toxins are expressed at every stage of development. The dual purpose in this case would be a consequence of the context-dependency of venom. In case of large host animals like cattle and horses the venom of adult tabanids mainly causes, immunosuppression, vasodilatation and prevents blood clotting all promoting blood feeding. The same venom though injected into a thousand times smaller prey organism could potentially evoke much more deleterious effects. It is noteworthy that the functional characterization of venom from adult tabanids was completely focused on the blood-feeding context. Hence potential insecticidal or cytotoxic effects have not been tested yet. An argument against a potential dual function of single venom compounds is that effects e. g. on the coagulation cascade would only affect vertebrate prey as invertebrates (the main prey of larval tabanids) lack a comparable coagulation cascade.

The second hypothesis is that tabanids possess two highly deviating venoms, one used for predation and the other for blood feeding. This would imply that during metamorphosis an expression shift of venom compounds takes place. Due to the high metabolic costs of venom such ontogenetic shifts in venom composition have frequently been observed, especially in the context of a shifted prey preference (Surm and Moran, 2021b). The functional annotation of venom precursors identified in the venom of larval stages of a tabanids provides a first idea regarding the main venom constituents utilized by larval tabanids to overpower prey. Especially the high diversity of discovered peptide toxins comprising putative neurotoxins and membrane-disrupting peptides indicates that cytotoxic effects combined with paralysis caused by neurotoxins are the main drivers of prey incapacitation in case of tabanid larvae.

The main constituents from adult tabanid venom that act on hemostatic processes are protease inhibitors. Likewise, also the larval venom contains several protease inhibitors and some of these show high similarity to Vasotab which inhibits vasoconstriction (Takáč et al., 2006b) or Tabkunin which mainly inhibits Thrombin (Xu et al., 2008) thereby interfering with the coagulation cascade. Interestingly none of these compounds are up-regulated in the larval venom as the expression levels are relatively low, especially compared to the highly expressed peptide toxins that were identified. This is a first indication for an expression shift of the venom compounds between tabanid life stages, although a direct comparison of the venom compound expression profiles within a single tabanid species is required to fully confirm this. It is noteworthy that a similar compositional shift was described for some rattle snakes. In this case the venom of the juveniles that mainly feed on lizards is more enriched with neurotoxins, whereas the venom of the adult snakes comprises more proteases and hemotoxic compounds utilized for envenoming mammals (Mackessy, 1988).

## 6. Acknowledgements

We thank the CECAD Proteomics Facility (University of Cologne) for excellent support with Quadrupole Orbitrap analyses. Further thanks go to the CCG (Cologne Center for Genomics) for providing high-quality transcriptome sequencing results. Finally, we would like to thank Reinhard Predel for promoting the preliminary work.

## 7. Author contributions (CRediT)

**Jonas Krämer**: Writing – review & editing, Writing – original draft, Visualization, Project administration, Methodology, Investigation, Formal analysis, Data curation, Conceptualization; **Ludwig Dersch**: Writing –review & editing, Investigation; **Andreas Vilcinskas**: Writing – review & editing, Supervision, Project administration; **Tim Lüddecke**: Writing – review & editing, Supervision, Project administration, Funding acquisition, Conceptualization

